# Towards Simulation Optimization: An Examination of the Impact of Scaling on Coalescent and Forward Simulations

**DOI:** 10.1101/2024.04.27.591463

**Authors:** Tessa Ferrari, Siyuan Feng, Xinjun Zhang, Jazlyn Mooney

**Affiliations:** Department of Quantitative and Computational Biology; University of Southern California, Los Angeles, CA, USA; Laboratory of Genetics, University of Wisconsin, Madison, WI, USA; Department of Human Genetics; University of Michigan, Ann Arbor, MI, USA

## Abstract

Scaling is a common practice in population genetic simulations to increase computational efficiency. However, there exists a dearth of standardized guidelines for best practices. Few studies have examined the effects of scaling on diversity and whether the results are directly comparable to unscaled and empirical data. We examine the effects of scaling in two model populations, modern humans and *Drosophila melanogaster*. The reason is twofold: 1) due to the substantial difference in population sizes and generation times, human populations require moderate-to-no scaling, while more dramatic scaling is required for *Drosophila*; and 2) model populations have empirical data for comparison. We determine whether coalescence, runtime, memory, estimates of diversity, the site frequency spectra, and deleterious variation are affected by scaling. We also explore the effect of varying the simulated segment length and burn-in times. We find that the typical 10N generation burn-in is often not sufficient for full coalescence to occur in human or *Drosophila* simulations. As expected, memory and runtime increase as the scaling coefficient decreases and the length of the simulated segment increases. We show that simulating larger segments in humans is preferable, as it produces a smaller variance in diversity estimates. Conversely, in *Drosophila* it is preferable to simulate smaller segments and concatenate them into full genome for achieving comparable levels of diversity to empirical data. We find that aggressive scaling leads to stronger negative selection and ultimately amplifies the strength of background selection on flanking variation.

**Author Summary:** Scaling is a common approach to make population genetic simulations more computationally tractable. However, the implications of scaling and best practices for scaling are still unknown. This study highlights the importance of carefully considering scaling practices for forward-in-time population genetics simulations. We provide insights about the trade-offs between computational efficiency and accuracy of scaled simulations relative to empirical data, in human and *Drosophila*. We achieved this by varying the species demographic model; the method of coalescence; the simulated genomic element length; and the scaling factor. For each combination of parameters genetic diversity was quantified and computational was efficiency tracked. Our findings suggest that when simulating populations, such as humans, where moderate scaling is required, one should simulate larger genomic segments for more accurate measures of diversity. Scaling seems to cause an inflation of diversity in human simulations relative to empirical data. On the other hand, in populations where more aggressive scaling is required, such as *Drosophila*, simulating smaller segments is advantageous. The scaling factor increases substantially in *Drosophila* studies, and the simulated data experiences a drastic drop in diversity, relative to empirical data, and an increased effect of purifying selection.

## Introduction

A fundamental aspect of population genetics studies is to understand empirical genetic data and make inferences about the evolutionary past. Accurate understanding of empirical data is often challenging, as the observed genetic variation patterns can result from various combinatorial scenarios of evolutionary processes and are only limitedly informative about the spatiotemporal dynamics of evolution. In the light of this, the rapid development of computational simulation tools, especially Forward-in-time simulation methods [1–5] such as SLiM [6], has begun to address these challenges. By modeling different evolutionary processes such as mutations, natural selection and demographic changes, simulations can powerfully bridge the gaps left by empirical data by capturing the full range of genetic diversity generated by various combinations of evolutionary parameters. Numerous breakthroughs have been enabled by SLiM-type simulations over the recent years, including but not limited to inferring demographic history [7–12], detecting adaptation events [13–18], deciphering complex trait genetics [18–21], conservation genomics [22–25], and allowing hypothesis testing and methods development [26–30] in population genetics.

Despite the inarguable power and wide usability of simulation tools, there are many feasibility challenges with large-scale simulations, mainly due to the constraints with computational resources. To bypass the computational limitations, a common practice used by studies is to “scale down” [31] their simulation parameters by a certain factor. This practice theoretically allows certain metrics, such as levels of diversity and site frequency spectra (SFS), to remain constant while decreasing computational complexity. For example, the evolution of 1,000 generations in a population of 10,000 individuals, with mutation rate 2e^-8^ (bp/generation) and recombination rate of 1e^-7^ (crossovers/bp), is equivalent to evolving 100 generations of 1,000 individuals, at mutation rate 2e^-7^ (bp/generation) and mutation rate 1e^-6^ (crossovers/bp) (see *Methods*). Furthermore, a critical step of forward-in-time simulation is the inclusion of a “burn-in” period prior to the beginning of the simulated demographic model, during which time, genetic diversity in the initial population establishes equilibrium through mutation-selection-migration balance [32,33]. A typical burn-in time is suggested to be 10xN generations, where N is the initial effective population size. Therefore, scaling down of simulations also implies a substantially reduced burn-in time, hence greatly improved computational efficiency.

However, regardless of its convenience, the practice of scaling is not without controversy [34,35]. The modification of fundamental evolutionary parameters can introduce biases and potential discrepancies between what is simulated and what is intended by the model [26,35–37]. As such, scaled simulations could ultimately lead to inaccurate downstream inference results that rely heavily on comparing scaled simulation data to empirical observations. Such concern is especially pronounced given there is a general lack of understanding of the aspects of, and the extent to which, the simulated data are compromised upon scaling. Because of this, we currently do not have standardized guidelines on how to appropriately scale simulations while minimizing these distortions.

This study aims to address this outstanding problem by investigating the performance and outcomes of scaling on SLiM-based simulations by comparing simulation results to empirical genetic data. We evaluate the scaling effects by quantifying several standard measures of genetic diversity, such as heterozygosity, Watterson’s theta, and the SFS, as well as the operational metrics for computational efficiency, including runtime and memory usage (Figure 1). Additionally, we examine the impact of scaling on the dynamics of deleterious variation and the coalescence during the burn-in period.

**Figure 1.**
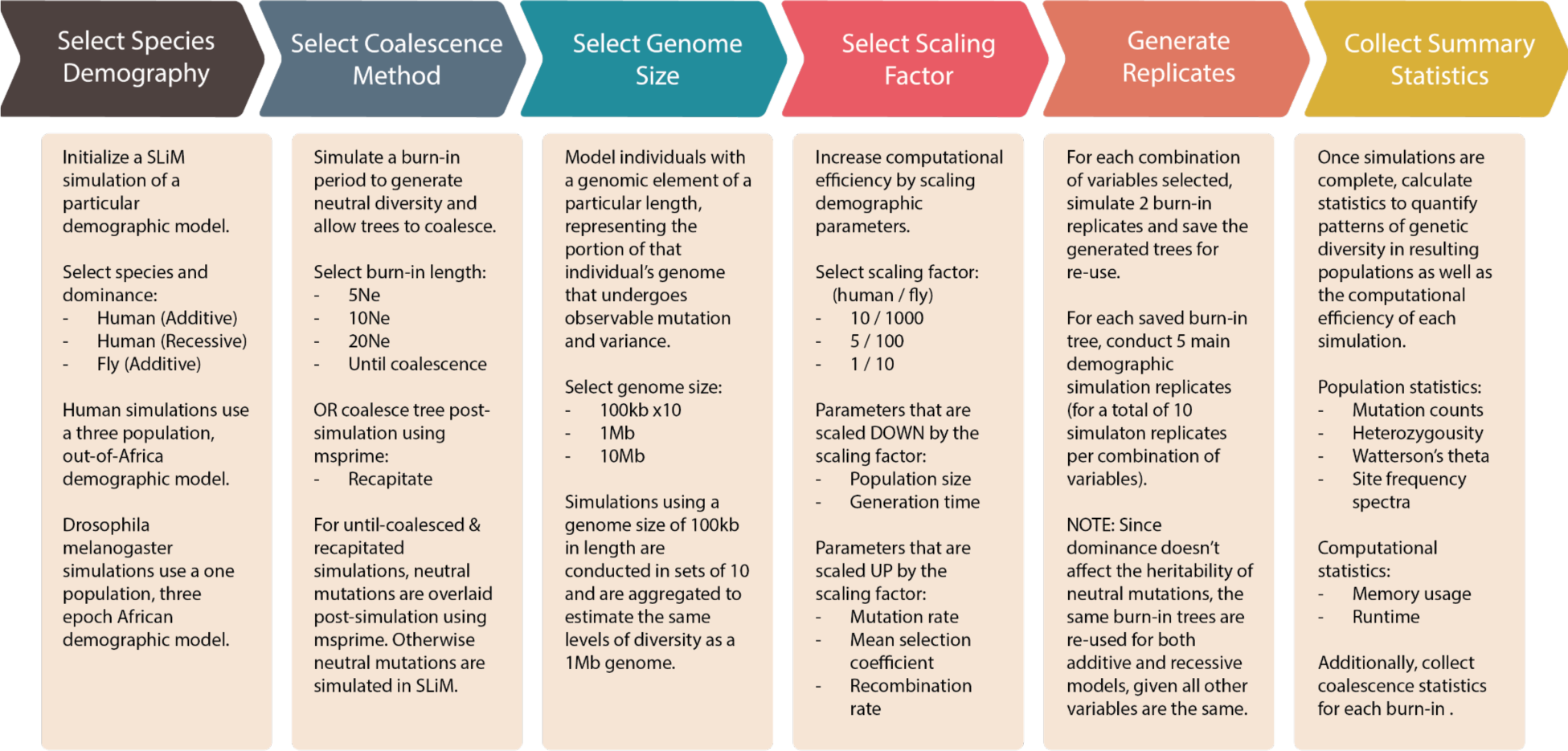
Summary of simulation parameters and workflow. Broadly, we conducted forward and coalescent simulations with two species (human and Drosophila) and varied genome size, scaling factor, and dominance (in the case of humans). We explored a total of 135 combinations of simulation parameters, where we conducted two burn-in replicates, except for simulations that were recapitated. The burn-ins were simulated with neutral dynamics, and we ran 5 main simulation replicates using each saved burn-in state, for a total of 10 replicates per parameter combination. Recapitated simulations were similarly conducted with 10 replicates per parameter combination. We additionally varied genome size, resulting in a grand total of 5,400 simulations across humans and flies.

Our work utilizes two classic study systems in population genetics: modern humans [38] and *Drosophila melanogaster* [39]. Given the inherent differences in population and life-history traits in the two species, we can use these two study systems to examine the effects of different magnitudes of scaling factors. In human studies, a scaling factor between 2-10 is generally sufficient. In *Drosophila*, populations require much more aggressive scaling such as a factor of 100-1000 to achieve reasonable computational usage. Additionally, both systems are characterized by extensive empirical data [40–48], which allows for robust comparisons of scaling performance.

By comprehensively evaluating scaling in these two model species, our work provides insight about how different scaling strategies can lead to more accurate and efficient simulation outcomes and allows for the development of guidelines of best practices for future population genetic application simulations.

## Results

### Simulation pipeline and burn-in coalescence

A common practice in SLiM is to simulate a burn-in period, which is intended to allow neutral diversity to reach an equilibrium and allow genealogical trees of all genetic loci to coalesce. A heuristic of 10N generations is often cited as a comprehensive standard for burn-in length. We simulated 18 total burn-ins, for each of the lengths 5N, 10N, and 20N (**Figure 1**) in each species. For both humans and flies, we found that none of the burn-ins coalesced with 5N. Importantly, for both flies and humans, 10N was also often not sufficient for full coalescence **(Table 1)**, no matter what scaling factor or genome length was simulated. Only four of the 18 total burn-ins with 10N reached full coalescence for humans (**Table 1A**) and none of the 10N *Drosophila* burn-ins fully coalesced (**Table 1B**). When we increased the burn-in length to 20N, most of the burn-ins coalesced for humans, however none of the *Drosophila* burn-ins coalesced.

**Table 1.**
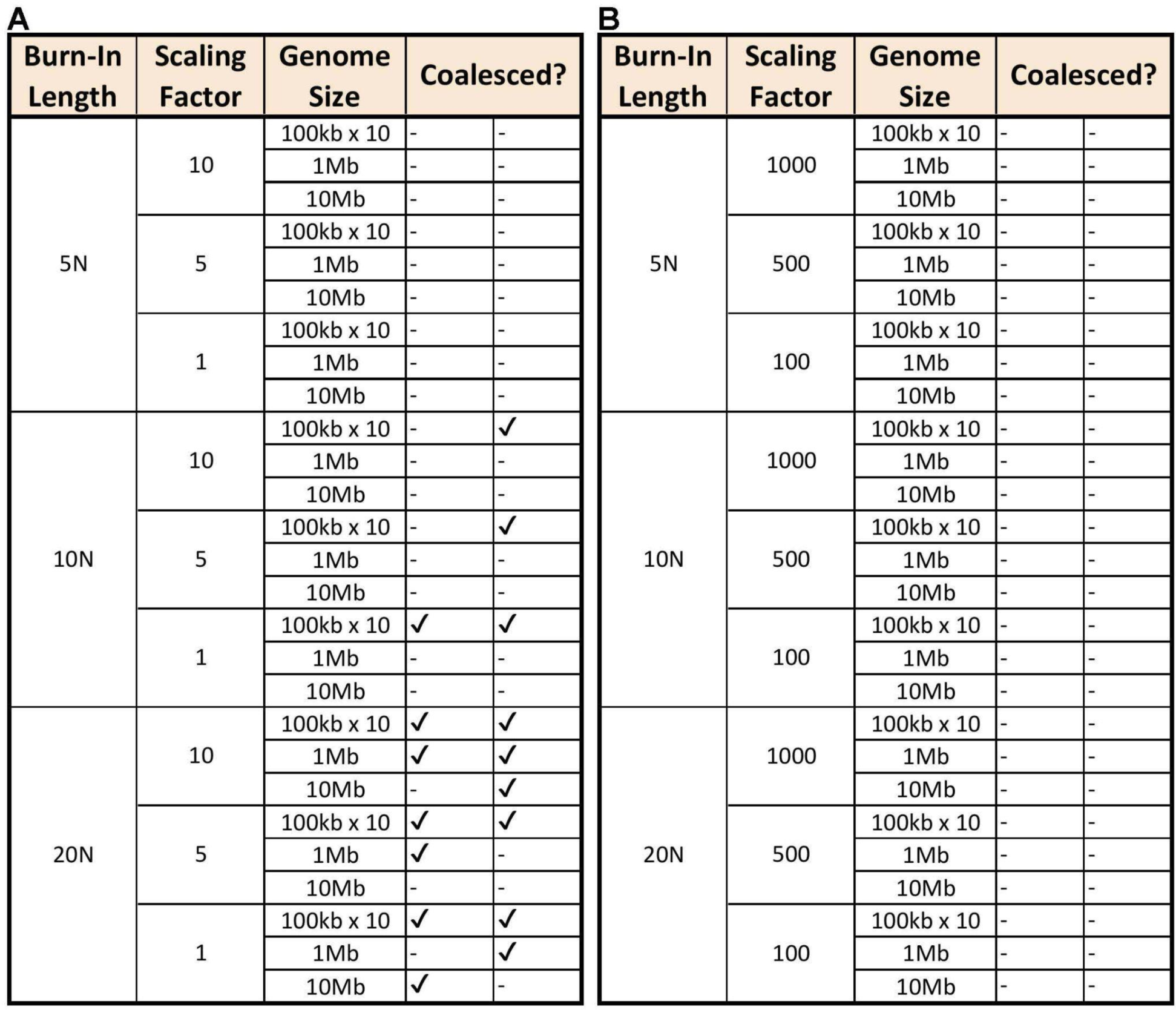
Coalescence of burn-in periods. Coalescence of A) human burn-in trees and B) Drosophila burn-in trees across parameter combinations. A minus sign (-) indicates that the burn-in replicate did not fully coalesce by the end of the burn-in period, while a check mark indicates that it did. About a quarter of the human burn-ins coalesced while none of the Drosophila replicates coalesced.

To explore the time to coalescence further, we ran additional simulations and used the *checkCoalesce* function in *msprime.* For these additional replicates, we observed an average time to coalescence of 17.57 ± 5.28N generations for human and 25.20 ± 3.98N generations for *Drosophila* (**Supplementary Table 1**). The fastest time to coalescence was 8.62N and the longest was 25.80N in humans (**Supplementary Table 1A**). Notably, for flies, we observed a minimum of 18.69N generations until coalescence, with a maximum that was greater than 30N generations (**Supplementary Table 1B**). We capped simulations at 30N for computational efficiency and since it is far beyond the common burn-in recommendation of 10N.

### Computational efficiency

We computed the total runtime of each simulation as the sum of the runtime of the coalescence method (neutral burn-in or recapitation), and the runtime of the main simulation (with neutral and deleterious mutations). Even with more dramatic scaling, non-recapitated *Drosophila* simulations’ runtimes ranged from approximately 2-25 times longer than human simulations with comparable parameters (**Figure 2 & Supplementary Tables S2&3**). Interestingly, when recapitation was used we observed similar runtimes in human and *Drosophila* simulations with analogous parameters (**Supplementary Tables S2&3**). Irrespective of species or dominance model, we observed that runtime decreased as the scaling factor increased and the simulated segment length decreased (**Figure 2 and Supplementary Tables S2&3**). This result was expected given the utility of scaling. For humans, total runtimes were not significantly different between additive and recessive models (**Figure 2**).

**Figure 2.**
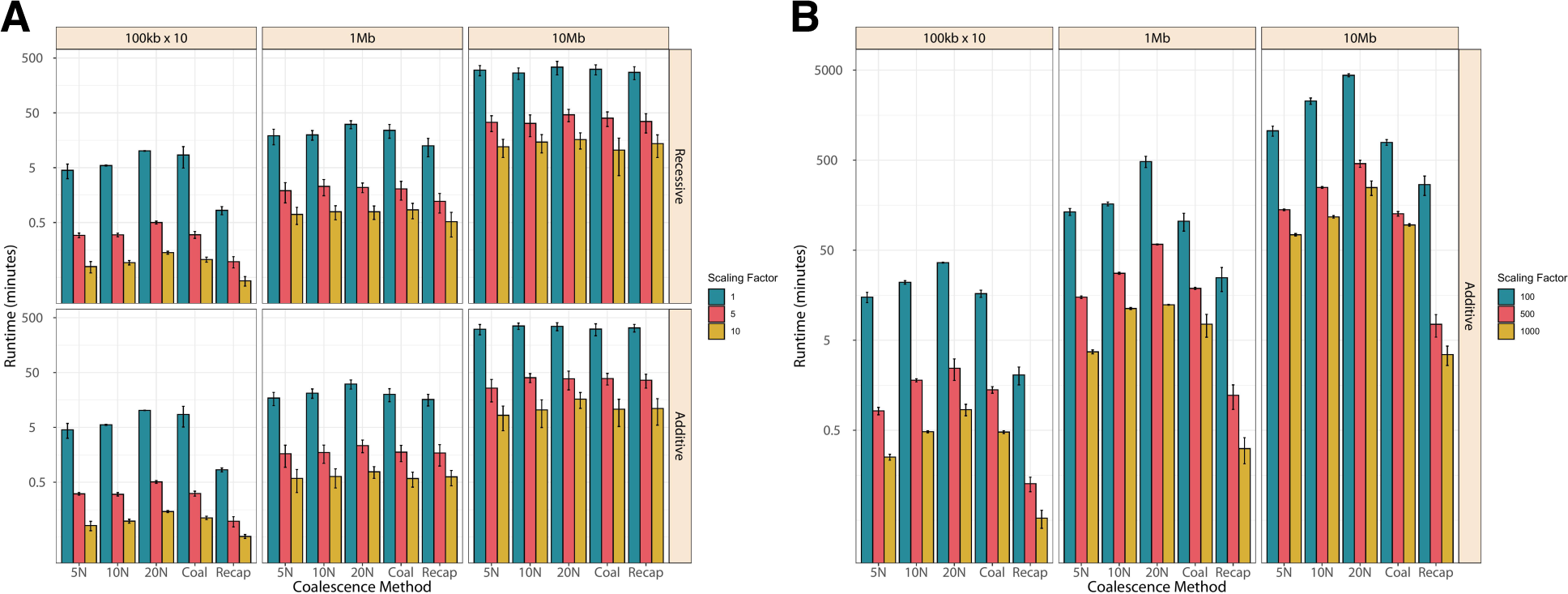
Total runtime of demographic simulations. Total runtime of A) human simulations and B) Drosophila simulations across 5 coalescent methods. For A) the top row used an additive model of dominance for mutations and the bottom row used a recessive model. Different colors represent different scaling factors, ranging from 1 (unscaled) to 10 in human or from 100 to 1000 in Drosophila. Each bar represents the average runtime of 10 simulation replicates with error bars showing standard deviation.

The fastest simulations were coalescent or recapitated irrespective of species (**Figure 2 and Supplementary Tables S2&3**). This was expected since neutral mutations are added post-hoc to trees, which will save the user time. Though SLiM coalescence checking and recapitation were the fastest coalescence methods for both human and *Drosophila* (**Supplementary Figure S1**), the computational gain for total simulation runtime became less apparent in humans as the simulated segments increased. This is because in humans, the coalescence method accounts for a smaller proportion of the total runtime, so when genome size is increased and the main simulation gets longer, differences in runtime due to the coalescence method become more negligible (**Figure 2 & Supplementary Figure S1A**). Interestingly, in *Drosophila*, recapitated simulations were always faster, with coalescent simulations being the second fastest (**Figure 2 and Supplementary Tables S3**). The computational efficiency of recapitation held across any scaling factor and the length of the simulated segment (**Supplementary Tables S2&3**).

We next quantified the peak memory usage of each set of simulation parameters (**Figure 3 and Supplementary Tables S2&3**). The peak memory is the maximum from the neutral burn-in and the main simulation with both neutral and deleterious variation. For both species, memory usage was higher in the main simulation than the burn-in for all parameter combinations (**Figure 3 and Supplementary Tables S2&S3**). The trend in runtime that we observed was precisely what we saw with computational time, where average memory usage decreased as the scaling factor increased and the simulated segment length decreased. At their height, 10 Megabases (Mb), human simulations peaked on average at 6,254 Megabytes (**Supplementary Tables S3**) and *Drosophila* simulations used approximately 4-5 times that amount and peaked at 29,556 Megabytes (**Supplementary Tables S3**).

**Figure 3.**
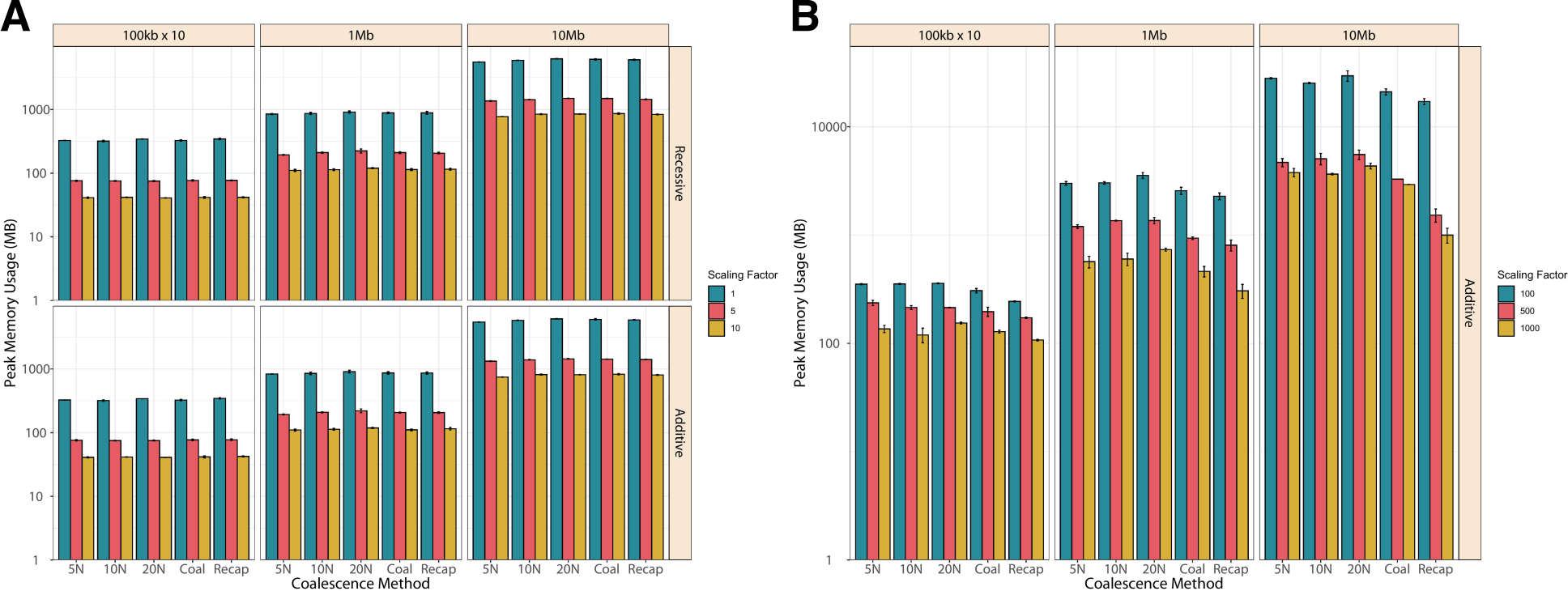
Peak memory usage of human demographic simulations. Peak memory usage of A) human simulations and B) Drosophila simulations across 5 coalescence methods. For A) the top row used an additive model of dominance for mutations and the bottom row used a recessive model. Different colors represent different scaling factors, ranging from 1 (unscaled) to 10 in human or from 100 to 1000 in Drosophila. Each bar represents the average peak memory usage of 10 simulation replicates with error bars showing standard deviation.

### Diversity in simulations versus empirical data

Next, we explored whether there were differences in genetic diversity between the simulated data relative to empirical data. We compared the simulated human African population to data from the 1000 Genome Yoruba (YRI) population [40] and compared the simulated *Drosophila* data to data from the Zambian DPGP3 population [46]. We generated an SFS for each population using exome data, which our simulations were designed to approximate, as well as the whole genome data.

For the human population (**Figure 4**), the SFS created from the simulated data was comparable to the empirical exome data across all parameter combinations. The largest variance in the SFS could be seen in the low frequency bins and simulations where the genomic length was smallest (100 Kilobases (Kb)). Genomic segments simulated to be 10Mb had the lowest standard deviation across all bins. The SFS generated using the whole genome sequence data (**Supplementary Figure S2**) was also quite comparable to the simulated data. However, in the simulated data there was a noticeable increase in the proportion of variants with MAF (0-0.05] relative to the empirical data.

**Figure 4.**
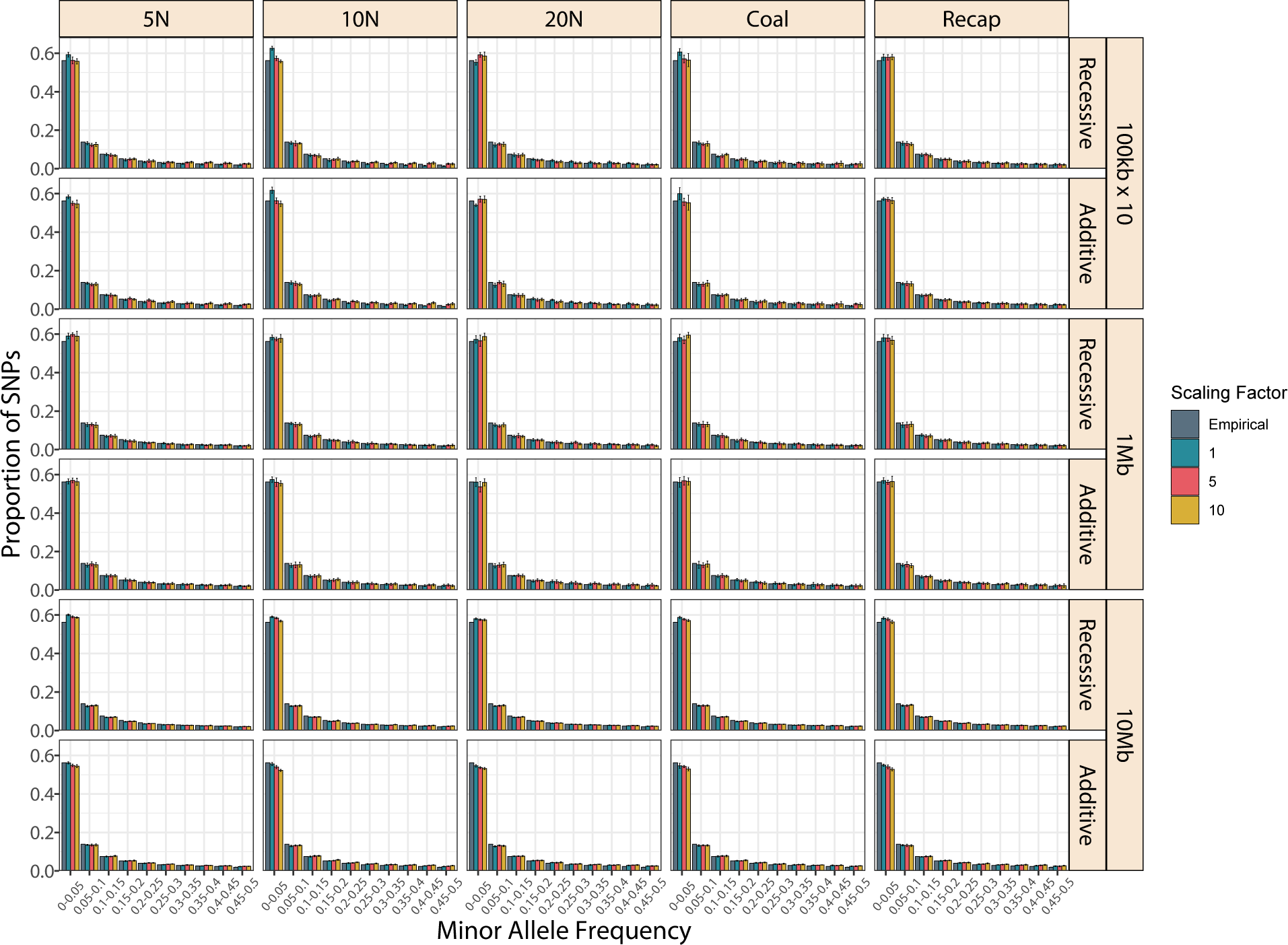
Exonic site frequency spectra for simulated and empirical Yoruban population. Each pair of rows represents a different simulated segment length, with either an additive or recessive model of dominance. Different colors represent different scaling factors, ranging from 1 (unscaled) to 10. Each bar represents the proportion of SNPs at a given MAF averaged over 10 simulation replicates, with error bars showing standard deviation. MAF bins are left-exclusive and right-inclusive, meaning fixed sites (MAF=0) are not included.

In *Drosophila* populations (**Figure 5**), the SFS created from simulated data had distributions that were skewed towards lower frequencies than those of the empirical exome data. This difference is especially evident in the leftmost (0,0.05] MAF bin, which contains the largest proportion of SNPs in the SFS. This pattern is magnified in the 10Mb simulations with the highest scaling factors (500 & 1000) where almost all the SNPs in the simulated data are skewed towards the lowest frequencies. The fraction of variants that are assigned to bins associated with more common variation (i.e. MAF > 0.1) within the empirical data overtook the simulated data across all parameter combinations. This same trend was observed in the whole genome sequence data (**Supplementary Figure S3**). In contrast to what we observed with the human data, the SFS created with the whole genome data more closely matched the simulations, because there were slightly more alleles in the lowest MAF bins. However, the difference between the fraction of sites in the lowest MAF bins in the whole genome and exome dataset is so small that they are qualitatively identical.

**Figure 5.**
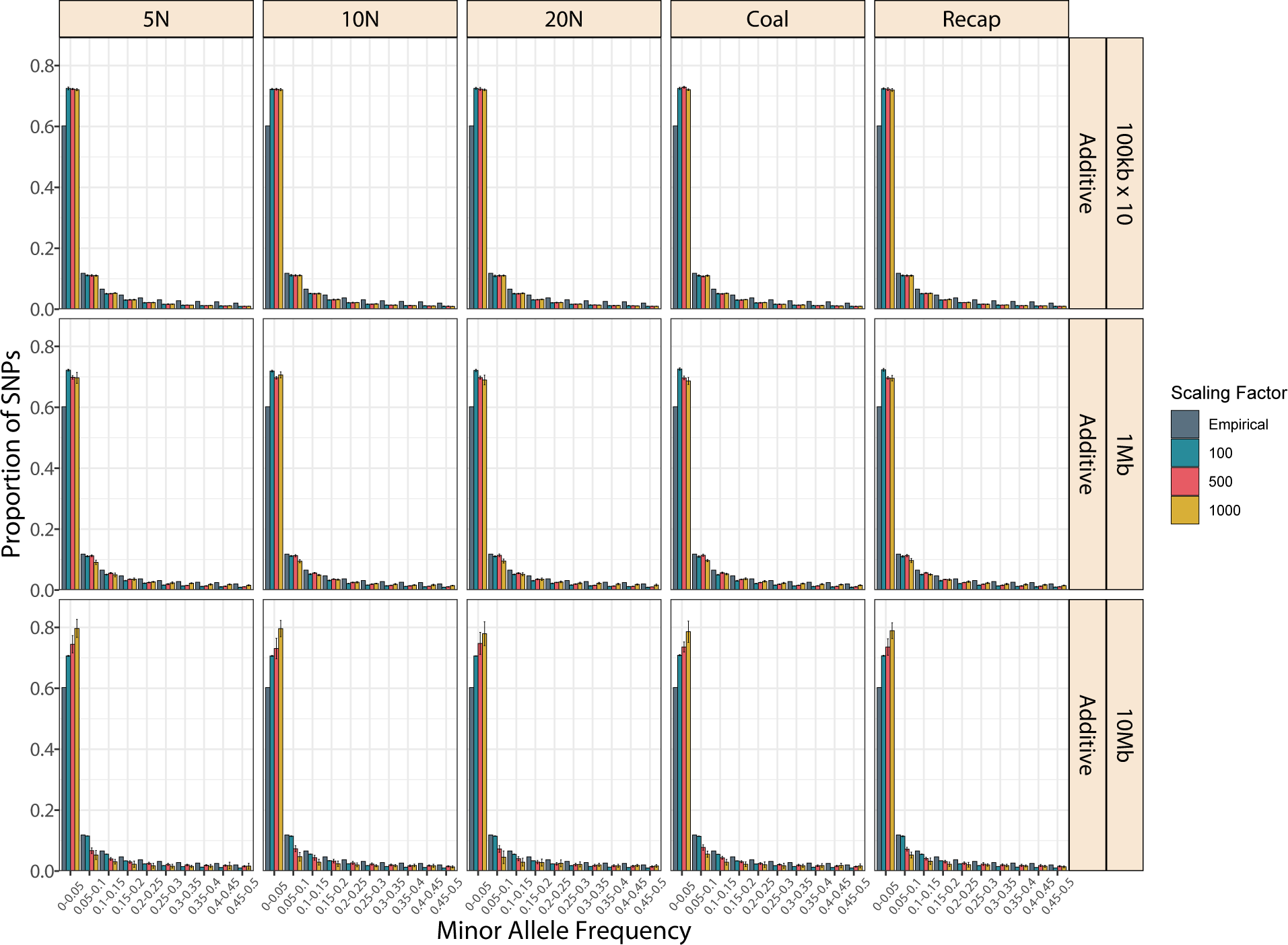
Exonic site frequency spectra for simulated and empirical African Drosophila population. Each row represents a different simulated segment length. Different colors represent different scaling factors, ranging from 100 to 1000. Each bar represents the proportion of SNPs at a given MAF averaged over 10 simulation replicates, with error bars showing standard deviation. MAF bins are left-exclusive and right-inclusive, meaning fixed sites (MAF=0) are not included.

Given that we observed differences between the SFS generated from simulated and empirical data for both human and *Drosophila* populations, we next decided to examine commonly used summaries of the SFS. We started by computing the heterozygosity from the SFS generated from exonic regions (**Figure 6**) and from the whole genome (**Supplementary Figure S4**). Strikingly, the data showed quite disparate patterns between human (**Figure 6A**) and *Drosophila* (**Figure 6B**).

**Figure 6.**
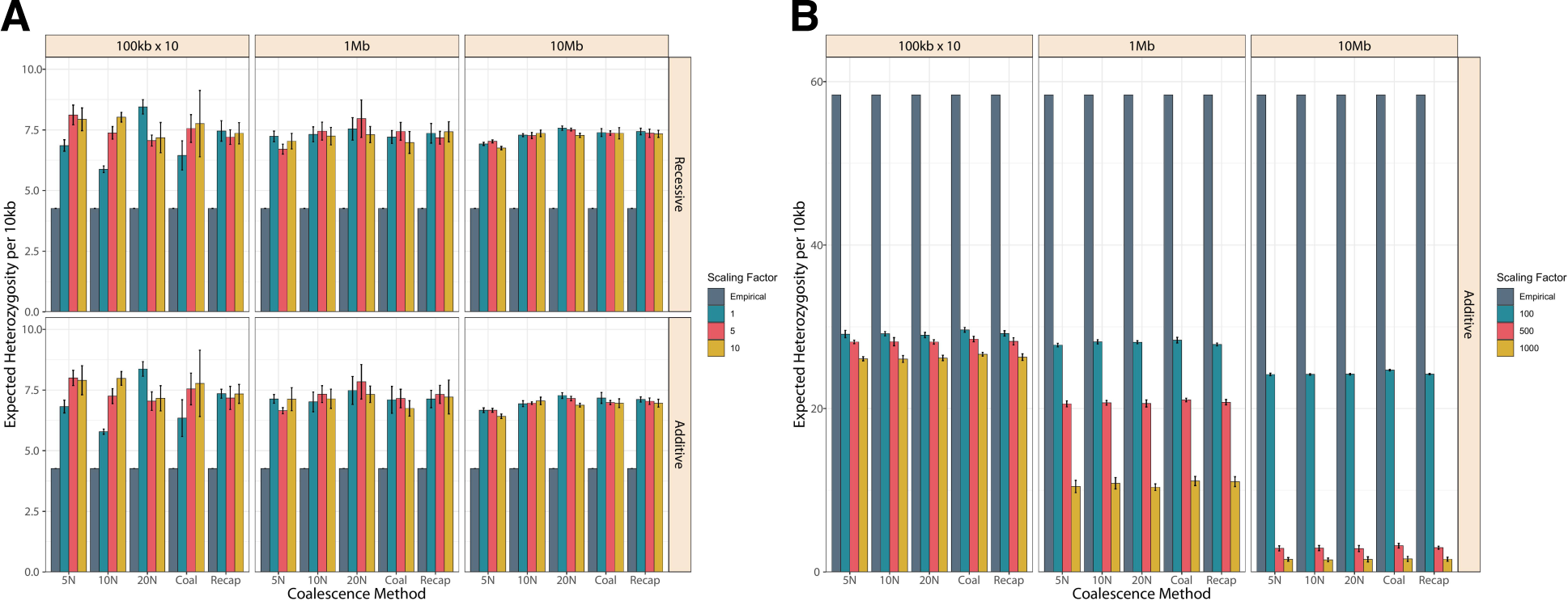
Heterozygosity computed from the exonic site frequency spectra for simulated and empirical African populations. A) For human data, the top row used an additive model of dominance for mutations and the bottom row used a recessive model. B) Drosophila data used only additive dominance. Different colors represent different scaling factors, ranging from 1 (unscaled) to 10 in humans or from 100 to 1000 in Drosophila. Each bar represents the average expected heterozygosity of 10 simulation replicates with error bars showing standard deviation.

For simulated human data, there was no significant difference between heterozygosity across simulation parameters, although simulations with smaller genome sizes generally had higher standard deviations (**Figure 6A & Supplementary Figure S4A**). Interestingly, we found that the heterozygosity computed from both exome data (**Figure 6A**) and whole genome data (**Supplementary Figure S4A**) was lower than the heterozygosity computed from any of our simulated data. However, despite simulations being designed to resemble exonic segments, the whole genome heterozygosity was much closer to simulated estimates than the exonic heterozygosity.

Conversely, for flies we observed a dramatic deficit in heterozygosity estimates from simulated data relative to both exome (**Figure 6B**) and whole genome (**Supplementary Figure S4B**) empirical data. This deficit was magnified in the whole genome data, with simulated estimates widely ranging from about 2.5-50 times lower than whole genome estimates. A greater deficit of expected heterozygosity was observed in simulations with larger scaling factors, and this effect was dramatically exacerbated if the simulated segment was longer. This deficit was stable across all coalescence methods. The simulations that most resembled empirical heterozygosity estimates were those with the lowest scaling factor (100) and the shortest segment length (100kb), however even these were underestimates.

Finally, we examined Watterson’s theta, which was also computed from the SFS generated using exonic regions (**Figure 7**) and the whole genome (**Supplementary Figure S5**). Largely, we observed the same pattern as with heterozygosity. The human (**Figure 7A**) and *Drosophila* (**Figure 7B**) populations had opposite outcomes when comparing the empirical versus simulated data.

**Figure 7.**
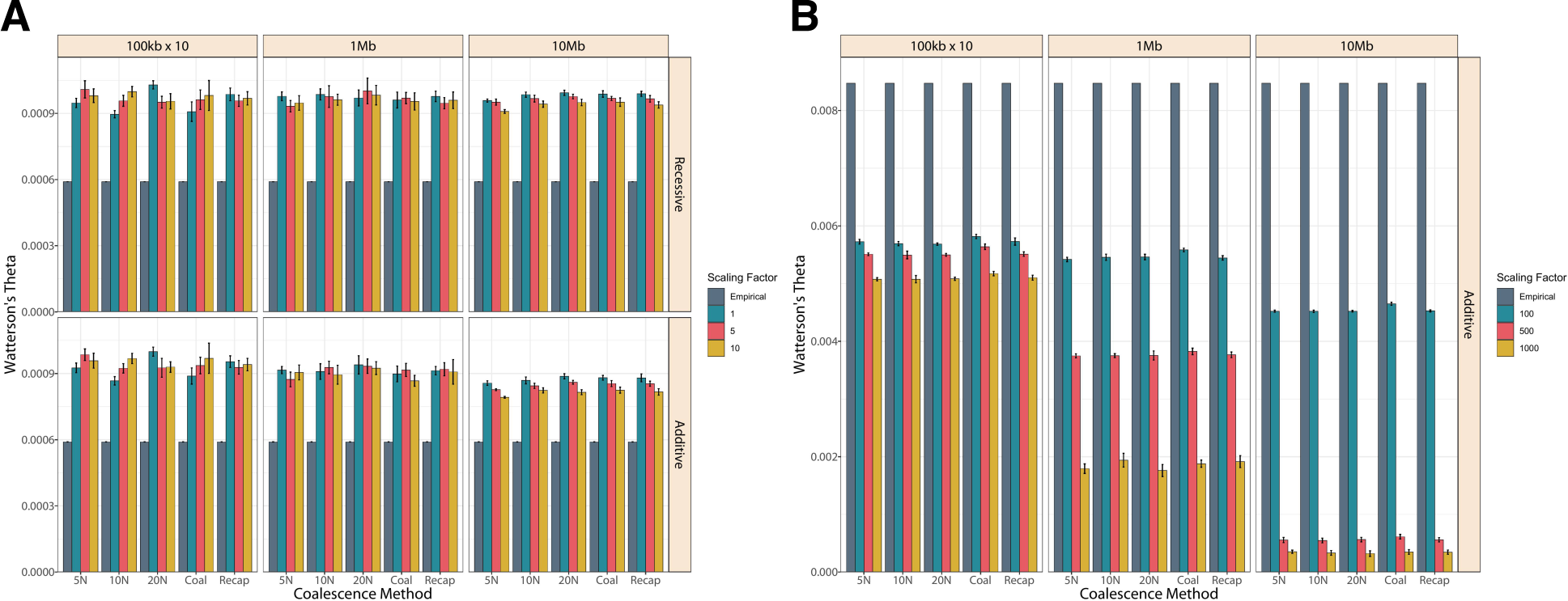
Watterson’s theta computed from the exonic site frequency spectra for simulated and empirical African populations. A) For human data, the top row used an additive model of dominance for mutations and the bottom row used a recessive model. B) Drosophila data used only additive dominance. Different colors represent different scaling factors, ranging from 1 (unscaled) to 10 in humans or from 100 to 1000 in Drosophila. Each bar represents the average value of Watterson’s theta for 10 simulation replicates with error bars showing standard deviation.

Once again, for the human data, there was no significant difference between theta across simulation parameters, and simulations with smaller genome sizes generally had higher standard deviations (**Figure 7A & Supplementary Figure S5A**). Theta computed from both exome data (**Figure 7A**) and whole genome data (**Supplementary Figure S5A**) was lower than theta computed from any of our simulated data. Whole genome theta estimates were much closer to simulated estimates than the exonic theta estimates.

For the flies, we observed a dramatic deficit in theta estimates from simulated data relative to both exome (**Figure 7B**) and whole genome (**Supplementary Figure S5B**) empirical data.

This deficit was magnified in the whole genome data, with simulated estimates widely ranging from about 2-35 times lower than whole genome estimates. A greater deficit of theta estimates was observed in simulations with larger scaling factors, and this effect was dramatically exacerbated if the simulated segment was longer. This deficit was stable across all coalescence methods. The simulations that most resembled empirical theta estimates were those with the lowest scaling factor (100) and the shortest segment length (100kb), however even these were underestimates.

## Discussion

By systematically comparing genetic diversity metrics, coalescent times, and computational efficiencies between two model organisms, our study highlights that those nuanced decisions on the choice of scaling factors can have major impacts on the accuracy of forward-in-time simulations in population genetics.

One of the most striking results was the outcome of coalescence in simulations at all scaling factors. Most simulations do not fully coalesce with burn-in periods less than 30N. We show that the heuristic of 10N generations for the burn-in period, which is often cited as a standard for burn-in length, is insufficient for all trees to fully coalesce in both human and *Drosophila* models, while it does allow neutral diversity to reach an equilibrium, this has been noted in the SLiM manual [49] and another recent study [34]. The lack of coalescence is particularly crucial as it highlights a potential area of oversight in simulations, where incomplete coalescence could lead to underestimation of linkage disequilibrium, and further affect the accuracy of haplotype-based population genetic inference methods. We recommend studies to re-evaluate burn-in lengths to ensure they are adequate for the specific dynamics of the population and scaling used, especially in studies where all lineages are required to coalesce. The SLiM program offers an option to periodically check for coalescence as the burn-in period progresses. This option, though increasing SLiM memory requirements by necessitating tree sequence recording, is ideal if desired simulation outputs are dependent on full tree coalescence. Tree sequence recording has the added benefit of allowing neutral mutations to be overlaid using *msprime* [50,51] after the simulation is complete, which is faster and more computationally efficient than simulating neutral mutations in SLiM. Coalescent simulations like *msprime* have the benefit of being much less computationally expensive but are unfortunately limited to simulating neutral dynamics. However, *msprime* recapitation enables the best of both worlds - simulating non-neutral dynamics with SLiM, followed by using *msprime* to retroactively connect trees with no burn-in period required.

We show another major impact of scaling, particularly with *Drosophila*, is a shift in the distribution of allele frequency spectrum caused by especially large scaling factors. In this case, large scaling factors means proportionally greater selective pressure in deleterious mutations, making it difficult for new mutations to persist longer than a few generations in a population. Consequently, this leads to a notable increase in the number of rare variants, especially rare variants like singletons and doubletons, relative to empirical data. We also observed a decrease in mutation ages as scaling factors increase (**Supplementary Figure S6**). This is especially the case when the simulated genomic segment is long, regardless of the scaling strength, as we note a flipped pattern in mutation age distribution between different scaling factors when segment lengths increase from <1Mb to multi-megabase level (**Supplementary Figure S6**). We believe the impact is exaggerated in the longer segments because any strongly deleterious mutation can accelerate the purging of linked mutations along with it, whereas simulating smaller segments can diffuse the strength of purging in regions flanking deleterious mutations.

Additionally, in *Drosophila* we observed a marked decrease in genetic diversity and allele frequencies as the scaling factor increased, with the effect being drastically amplified when longer segments were simulated. This is likely due to the higher selective pressures exerted by more aggressive scaling. However, this observation also suggests that the effects of aggressive scaling are best combated by simulating multiple shorter genomic segments and subsequently aggregating them to approximate larger ones. This strategy is particularly promising as it allows accurate simulation of populations which require aggressive scaling and may also enable full genome simulations in population genetics. The only limitation with concatenating smaller regions would be the level of LD. However, so long as the choices of small pieces are backed up by the empirical recombination map (meaning that the small segments are picked by recombination breaking points), the LD patterns in simulations should not be of concern.

Conversely, for humans where moderate scaling factors are used, we show that simulated SFS and empirical observations are highly concordant. In terms of the levels of genetic diversity, we show that heterozygosity and Watterson’s theta can be slightly overestimated in scaled simulations but generally comparable to empirical data. Additionally, larger simulated segments (multi-megabase) generally produced more stable estimates. This suggests that with moderate scaling, simulating longer genomic regions is preferred regardless of scaling factor, as this approach minimizes noise to capture more precise representations of genetic diversity.

Finally, as expected, we show that the memory usage and runtime increased with decreased scaling coefficients and longer simulated segments, indicating that the reduction in computational costs will remain the major motivation for scaling in population genetic simulations. This trade-off between computational demand and simulation accuracy emphasizes the importance of optimizing both scaling and computational resource allocation to achieve the best possible outcomes without compromising the reliability of simulation results.

Taking these findings into account, a generalized strategy for scaled simulations becomes apparent. For simulating coalescence we recommend studies to use *msprime* recapitation, as it is generally the most computationally efficient and effective method. This is especially advantageous when the runtime for the classical burn-ins is long, similar to what we observed in *Drosophila*. Additionally, we recommend studies utilizing scaled simulations to choose genomic segment lengths that are both long enough to minimize noise, and short enough to combat scaling effects, while using the lowest scaling factor that is computationally feasible. If a study wishes to approximate a genomic length larger than the determined optimum, then simulations can be conducted in sets of smaller segments and aggregated to achieve the desired genomic length. This is particularly important for the study of Drosophila-like populations where aggressive scaling is necessary. These recommendations are by no means meant to be universally applicable. Rather, they are intended to provide a basic framework for scaled simulations which should be tailored to the species of interest and to the needs of the study.

In conclusion, our study provides critical insights into the scaling practices in forward-in-time simulations and their implications for research in population genetics. By addressing these challenges and providing evidence-based recommendations, we aim to contribute towards more accurate, efficient, and reproducible simulation methodologies in the field. We urge future research to focus on developing a framework for determining optimal scaling strategies that minimize biases while maximizing computational efficiency. The differential impacts of scaling on human-like populations and *Drosophila*-like populations underline the importance of tailored simulation strategies. It becomes evident that there is no one-size-fits-all approach to scaling in population genetic simulations. Instead, our findings advocate for a species-specific customization of simulation parameters, which considers the unique biological and life-history traits of the organism being studied.

## Materials and Methods

### Simulation overview

Forward-in-time simulations were conducted using SLiM 4.0.1 [52] on a high-performance computing cluster, utilizing Dell node models (R6525, R440, xl170, C6420) with core speeds ranging from 2.1-2.6Ghz. For simulations utilizing recapitation and/or neutral mutation overlay, tree sequence data was augmented using python 3.12.2 with packages *msprime* 1.3.1 [50], pyslim 1.0.4, and tskit 0.5.6 [53,54].

The simulation parameters and framework are summarized in **Figure 1**. The parameters that were varied were: 1) the species demographic model; 2) the method of coalescence; 3) the genomic element length simulated per individual, sometimes referred to as genome size; and 4) the scaling factor. For each combination of parameters, simulation replicates were generated, and summary statistics were calculated to quantify population composition and computational efficiency.

### Species Demographic Model and Simulations

Simulations were conducted using either human or *Drosophila melanogaster* demographic models. Population sizes for human demographic models were low enough that unscaled simulations could be conducted in a tractable time with reasonable computational efficiency. However, due to *Drosophila* population sizes being quite large, simulations are computationally intractable unless scaling is employed, and in some cases, computationally inefficient unless scaling is dramatic.

Human simulations utilized a three population out-of-Africa model, with SLiM scripts modified from [55], which closely follows [38], with an ancestral African population expansion, an out-of-Africa population split, followed by the divergence and exponential growth of the European and East Asian subpopulations. Migration between subpopulations is present throughout the model. In unscaled human simulations, we imposed a mutation rate of 2.36e-8 [38], a neutral to deleterious mutation ratio of 9:1, selection coefficients of new mutations are drawn from a gamma distribution with a mean selection coefficient of -0.03, a shape parameter of 0.2 [29], and a recombination rate of 1e-8.

*Drosophila* simulations utilized a three epoch African population model, with demographic specifications from Sheehan and Song [39]. This model outlines a bottleneck of the ancestral African population, followed by a recent population expansion. In *Drosophila* simulations, we imposed a mutation rate of 8.4e-9 [56], a neutral to deleterious mutation ratio of 1:2.85, selection coefficients of new mutations drawn (DFE) from a gamma distribution with a mean selection coefficient of -1.33e-4, a shape parameter of 0.35 [57], and a recombination rate of 2.06e-8 [36].

Human simulations were further stratified by dominance type, either recessive or additive. *Drosophila* simulations were only conducted with additive dominance, due to the computational intractability of recessive simulations with a large population size.

We also evaluated the efficiency of three coalescent methods: 1) classical burn-in of length 5N, 10N, or 20N; 2) burn-in with SLiM coalescence checking (denoted as *Coal*); and 3) msprime recapitation (denoted as *Recap*). For the latter two methods, neutral mutations were overlaid using *msprime* after the main simulation was completed.

### Genome Size

We conducted simulations with varying genome sizes 100kb x10, 1Mb, and 10Mb, for comparison. Simulations with a genome size of 100kb x10 consisted of a set of ten 100kb simulations, and summary statistics were aggregated. This made the 100kb x 10 comparable to the 1Mb simulation. This allowed us to effectively parallelize a 1Mb genome using a scatter-gather approach.

Genomic elements in SLiM were simulated uniformly across the whole segment with the mutation rate, neutral to deleterious ratio, DFE, and recombination rate held constant across all loci. These genomic segments are therefore only intended to simulate a simplified version of exome data.

### Scaling Factor

Simulations were scaled according to procedures outlined in the SLiM manual, by a constant scaling factor, Q. When scaling SLiM simulations, population size and generation time were scaled *down* by a factor of Q. Conversely, mutation rate, mean selection coefficient, and recombination rate were scaled *up* by a factor of Q.

Simulations were stratified by the magnitude of their scaling factor, Q. Human simulations used scaling factors of 1 (unscaled), 5, and 10. For *Drosophila* simulations, we increased the values of Q a hundredfold, with scaling factors of 100, 500, and 1000.

### Replicates

Across each species’ demography, coalescence method, genome size, and scaling factor, there were a total of 135 combinations of simulation parameters possible. For each parameter combination (apart from those that used recapitation) we conducted 2 burn-in replicates. The burn-ins were simulated with purely neutral dynamics and were re-used for both additive and recessive simulations. Tree-sequence recording was activated to enable burn-in re-use. We simulated 144 burn-ins total, saving their resulting trees. Then, we conducted 5 replicates of the main simulation using the saved burn-in state as a starting point, for a total of 10 replicates per parameter combination. Recapitated simulations were similarly conducted with 10 replicates per parameter combination. Since recapitation is randomized based on the ending state of the main simulation, each recapitated replicate effectively had a unique starting state. When we factored in varied genome size, a grand total of 5,400 simulations were conducted.

### Additional Burn-In Evaluation

To supplement our findings and provide additional information on coalescence in SLiM, we evaluated the efficacy of each burn-in method by quantifying coalescence. For classical burn-ins with lengths of 5N, 10N, and 20N we determined whether genealogical trees at all loci were fully coalesced or not using *tskit*. For burn-ins utilizing SLiM coalescence checking, we recorded how many generations each burn-in took to coalesce, expressed in terms of N.

### Human Empirical Data

We used the Yoruba (YRI) population from the 1000 Genomes [40] as our representative African human population for empirical data. The site frequency spectra (SFS) were generated following the approach in [7]. Previously, the authors identified 30 unrelated individuals, and removed 16 SNPs with missing data, leaving a total of 57,597,196 SNPs to generate the whole genome SFS. The canonical transcripts for hg19 were downloaded from UCSC genome browser [58] and used to subset exonic regions from the whole genome sequence data. Exonic regions were downloaded using UCSC Table Browser [59].

### Drosophila Empirical Data

To obtain empirical data for *Drosophila melanogaster*, we downloaded VCF files from *Drosophila* Genome Nexus (https://www.johnpool.net/genomes.html), which include the genome-wide genotype information of 197 individuals in an ancestral population (Siavonga, Zambia) [46]. High-quality sites across major chromosomal arms 2R, 2L, 3R, 3L and X were extracted by applying quality filtering to the VCF files as indicated in Lack et al. 2015. We subsequently removed the X chromosome. Allele counts from the remaining sites were then filtered to only biallelic sites and used for estimation of the folded SFS and genetic diversity. To estimate SFS and genetic diversity across exonic regions, genomic coordinates of exons (*Drosophila melanogaster* Genome assembly Release 5) were downloaded from UCSC Table Browser [59] to extract exonic sites (https://genome.ucsc.edu/cgi-bin/hgTables).

### Summary Statistics

Peak memory usage and runtime were recorded to quantify computational efficiency. While conducting simulations in SLiM, these metrics were calculated by specifying the -mem and -time flags at runtime. For running commands in python with *msprime*, runtime was recorded using the python time module, and memory usage was not recorded (but is assumed to be almost negligible).

We used *msprime* for post-processing simulations, subsequently returning to SLiM to generate the final outputs. Mutation data outputted by SLiM was used to quantify mutation counts (neutral, deleterious, and total). We generated summary statistics for the site frequency spectra (SFS) using BCFTools 1.10.2 [60] and VCFTools 0.1.14 [61].

Heterozygosity and theta were computed from the SFS, which was created by sampling 50 diploid individuals from the simulated data and 30 from the empirical. Expected heterozygosity was calculated assuming Hardy-Weinberg equilibrium as the sum of 2pq over all segregating sites. Theta was computed by tallying the number of segregating sites and dividing by Watterson’s constant [62]. We then normalized diversity by dividing it by the total length of the simulated sequence (i.e., genome size) or by the length of the empirical region of interest.

Specifically, the denominators for calculating empirical statistics were 2,897,310,462bp and 26,824,075bp for human (hg19) whole genome and exonic regions, respectively. For *Drosophila* (dm3/R5) the denominators were 96,606,862bp and 28,512,719bp for the whole genome and exonic regions, respectively. The SFS for *Drosophila* was made by down sampling the haplotype information to N=30 diploid genomes, so that it was comparable to the human sample size.

## Data Availability

Simulation code, resulting data, and scripts for analysis and plotting are available at https://github.com/TessaFerrari/Scaling_SLiM, and summary files can be found on Dryad.

## Acknowledgments

The authors would like to thank John Pool for his assistance with procuring and generating the *Drosophila* allele count data. We would also like to acknowledge Bernard Kim for helpful conversations and code.

## Supplementary Figures

**Supplementary Table S1.** Coalescent Simulations. Time to coalescence for A) human simulations and B) *Drosophila* simulations with coalescence checking in SLiM, using the checkCoalescence parameter.

| Burn-In Method | Scaling Factor | Genome Size | Coalesced? |  |
| --- | --- | --- | --- | --- |
| Coal | 10 | 100kb x 10 | 12.58N | 14.20N |
|  |  | 1Mb | 21.41N | 14.88N |
|  |  | 10Mb | 22.78N | 25.80N |
|  | 5 | 100kb x 10 | 11.94N | 8.86N |
|  |  | 1Mb | 18.35N | 16.48N |
|  |  | 10Mb | 20.92N | 24.83N |
|  | 1 | 100kb x 10 | 8.62N | 22.09N |
|  |  | 1Mb | 17.21N | 13.27N |
|  |  | 10Mb | 22.62N | 19.50N |

| Burn-In Method | Scaling Factor | Genome Size | Coalesced? |  |
| --- | --- | --- | --- | --- |
| Coal | 1000 | 100kb x 10 | 21.48N | 21.48N |
|  |  | 1Mb | 30.00N | - |
|  |  | 10Mb | - | - |
|  | 500 | 100kb x 10 | 23.99N | 28.52N |
|  |  | 1Mb | 29.41N | 27.57N |
|  |  | 10Mb | - | - |
|  | 100 | 100kb x 10 | 18.69N | 22.77N |
|  |  | 1Mb | - | 28.06N |
|  |  | 10Mb | - | - |

**Supplementary Table S2.** Human simulation computational summary statistics. Runtime and Peak memory usage for all human simulations across genome size, scaling factors, and dominance models.

| Burn-In Length | Scaling Factor | Genome Size | Runtime (minutes) |  |  |  | Peak Memory Usage (MB) |  |  |  |
| --- | --- | --- | --- | --- | --- | --- | --- | --- | --- | --- |
|  |  |  | Additive |  | Recessive |  | Additive |  | Recessive |  |
|  |  |  | Mean | SD | Mean | SD | Mean | SD | Mean | SD |
| 5N | 10 | 100kb x 10 | 0.08 | 0.02 | 0.08 | 0.02 | 41.00 | 0.87 | 41.26 | 0.97 |
|  |  | 1Mb | 0.59 | 0.26 | 0.70 | 0.24 | 110.08 | 4.09 | 110.58 | 4.43 |
|  |  | 10Mb | 8.31 | 3.96 | 12.05 | 4.39 | 747.44 | 5.25 | 775.66 | 3.18 |
|  | 5 | 100kb x 10 | 0.31 | 0.02 | 0.29 | 0.03 | 75.96 | 1.62 | 76.04 | 1.55 |
|  |  | 1Mb | 1.66 | 0.72 | 1.91 | 0.77 | 193.67 | 3.20 | 193.81 | 3.35 |
|  |  | 10Mb | 26.04 | 11.44 | 33.57 | 10.76 | 1327.44 | 17.48 | 1364.11 | 21.57 |
|  | 1 | 100kb x 10 | 4.54 | 1.37 | 4.48 | 1.33 | 326.50 | 2.38 | 326.07 | 2.88 |
|  |  | 1Mb | 17.14 | 4.62 | 19.11 | 6.00 | 836.92 | 6.01 | 848.37 | 19.28 |
|  |  | 10Mb | 309.43 | 65.55 | 300.27 | 62.91 | 5455.48 | 35.03 | 5561.57 | 47.30 |
| 10N | 10 | 100kb x 10 | 0.10 | 0.01 | 0.09 | 0.01 | 41.56 | 0.39 | 41.74 | 0.43 |
|  |  | 1Mb | 0.64 | 0.24 | 0.79 | 0.23 | 113.28 | 4.00 | 113.34 | 3.48 |
|  |  | 10Mb | 10.45 | 5.47 | 14.65 | 5.35 | 818.80 | 14.74 | 845.80 | 14.51 |
|  | 5 | 100kb x 10 | 0.30 | 0.02 | 0.30 | 0.02 | 75.14 | 1.05 | 75.51 | 1.40 |
|  |  | 1Mb | 1.74 | 0.64 | 2.29 | 0.75 | 208.89 | 4.20 | 210.35 | 5.25 |
|  |  | 10Mb | 40.40 | 7.51 | 32.05 | 13.69 | 1395.38 | 14.30 | 1431.19 | 8.67 |
|  | 1 | 100kb x 10 | 5.59 | 0.13 | 5.46 | 0.14 | 319.92 | 7.49 | 318.81 | 8.41 |
|  |  | 1Mb | 21.09 | 4.12 | 19.84 | 3.94 | 855.10 | 38.73 | 864.29 | 35.67 |
|  |  | 10Mb | 353.30 | 46.64 | 265.80 | 62.42 | 5810.50 | 69.64 | 5922.15 | 57.71 |
| 20N | 10 | 100kb x 10 | 0.15 | 0.01 | 0.14 | 0.01 | 40.90 | 0.30 | 40.86 | 0.36 |
|  |  | 1Mb | 0.78 | 0.19 | 0.78 | 0.21 | 118.91 | 2.34 | 120.18 | 2.52 |
|  |  | 10Mb | 16.37 | 5.23 | 16.27 | 5.33 | 815.56 | 5.93 | 847.02 | 7.34 |
|  | 5 | 100kb x 10 | 0.51 | 0.03 | 0.50 | 0.03 | 75.10 | 1.15 | 75.20 | 1.17 |
|  |  | 1Mb | 2.34 | 0.60 | 2.18 | 0.44 | 220.88 | 15.02 | 223.49 | 16.49 |
|  |  | 10Mb | 38.64 | 14.46 | 46.11 | 11.68 | 1444.31 | 18.56 | 1493.96 | 10.53 |
|  | 1 | 100kb x 10 | 10.22 | 0.06 | 10.13 | 0.15 | 341.96 | 1.51 | 343.02 | 2.09 |
|  |  | 1Mb | 30.83 | 5.80 | 30.77 | 5.08 | 911.96 | 41.70 | 912.49 | 35.66 |
|  |  | 10Mb | 347.81 | 58.14 | 339.96 | 94.04 | 6132.73 | 75.03 | 6253.77 | 63.77 |
| Coal | 10 | 100kb x 10 | 0.11 | 0.01 | 0.11 | 0.01 | 41.68 | 1.57 | 41.64 | 1.29 |
|  |  | 1Mb | 0.59 | 0.18 | 0.85 | 0.27 | 110.61 | 3.76 | 113.70 | 4.01 |
|  |  | 10Mb | 10.76 | 5.56 | 10.43 | 6.85 | 828.00 | 18.79 | 862.33 | 22.82 |
|  | 5 | 100kb x 10 | 0.31 | 0.03 | 0.30 | 0.04 | 77.01 | 2.01 | 76.89 | 2.08 |
|  |  | 1Mb | 1.77 | 0.58 | 2.05 | 0.77 | 207.20 | 3.88 | 210.38 | 6.94 |
|  |  | 10Mb | 39.02 | 9.25 | 40.06 | 11.95 | 1430.13 | 11.73 | 1485.67 | 14.25 |
|  | 1 | 100kb x 10 | 8.62 | 3.52 | 8.51 | 3.58 | 325.74 | 8.08 | 326.24 | 8.42 |
|  |  | 1Mb | 20.10 | 5.34 | 23.93 | 6.69 | 869.36 | 38.76 | 883.82 | 23.14 |
|  |  | 10Mb | 310.76 | 75.97 | 310.77 | 64.75 | 6016.90 | 171.52 | 6140.55 | 140.30 |
| Recap | 10 | 100kb x 10 | 0.05 | 0.00 | 0.04 | 0.01 | 42.44 | 0.79 | 41.73 | 0.68 |
|  |  | 1Mb | 0.63 | 0.20 | 0.52 | 0.25 | 115.33 | 5.26 | 115.57 | 3.83 |
|  |  | 10Mb | 11.11 | 5.61 | 13.74 | 6.09 | 805.93 | 12.63 | 835.45 | 13.12 |
|  | 5 | 100kb x 10 | 0.10 | 0.02 | 0.10 | 0.02 | 77.51 | 2.12 | 77.07 | 1.17 |
|  |  | 1Mb | 1.71 | 0.72 | 1.22 | 0.48 | 206.76 | 5.95 | 206.41 | 6.03 |
|  |  | 10Mb | 36.38 | 10.30 | 34.76 | 13.35 | 1407.82 | 13.99 | 1443.80 | 28.39 |
|  | 1 | 100kb x 10 | 0.84 | 0.07 | 0.83 | 0.14 | 345.68 | 7.99 | 344.96 | 11.25 |
|  |  | 1Mb | 16.16 | 3.87 | 12.52 | 4.62 | 864.93 | 33.30 | 884.52 | 47.42 |
|  |  | 10Mb | 326.51 | 51.28 | 273.45 | 73.03 | 5912.01 | 70.42 | 6041.01 | 111.33 |

**Supplementary Table S3.** *Drosophila* simulation computational summary statistics. Runtime and Peak memory usage for all human simulations across genome size and scaling factors.

| Burn-In Length | Scaling Factor | Genome Size | Runtime (minutes) |  | Peak Memory Usage (MB) |  |
| --- | --- | --- | --- | --- | --- | --- |
|  |  |  | Additive |  | Additive |  |
|  |  |  | Mean | SD | Mean | SD |
| 5N | 1000 | 100kb x 10 | 0.25 | 0.02 | 136.18 | 10.32 |
|  |  | 1Mb | 3.72 | 0.18 | 567.72 | 70.23 |
|  |  | 10Mb | 74.35 | 2.29 | 3781.83 | 336.17 |
|  | 500 | 100kb x 10 | 0.82 | 0.08 | 236.86 | 13.18 |
|  |  | 1Mb | 15.07 | 0.49 | 1199.20 | 47.76 |
|  |  | 10Mb | 140.82 | 2.76 | 4686.03 | 403.84 |
|  | 100 | 100kb x 10 | 15.08 | 1.98 | 352.31 | 4.11 |
|  |  | 1Mb | 133.39 | 11.74 | 3007.88 | 112.92 |
|  |  | 10Mb | 1058.87 | 134.61 | 28070.87 | 384.99 |
| 10N | 1000 | 100kb x 10 | 0.48 | 0.01 | 119.69 | 18.39 |
|  |  | 1Mb | 11.28 | 0.31 | 601.41 | 78.43 |
|  |  | 10Mb | 117.39 | 3.82 | 3657.38 | 69.42 |
|  | 500 | 100kb x 10 | 1.80 | 0.07 | 214.32 | 8.87 |
|  |  | 1Mb | 27.69 | 0.68 | 1359.62 | 14.65 |
|  |  | 10Mb | 249.02 | 6.30 | 5064.42 | 607.44 |
|  | 100 | 100kb x 10 | 22.01 | 1.02 | 354.10 | 3.34 |
|  |  | 1Mb | 163.46 | 8.10 | 3036.91 | 80.73 |
|  |  | 10Mb | 2269.55 | 183.24 | 25292.68 | 384.58 |
| 20N | 1000 | 100kb x 10 | 0.85 | 0.12 | 154.20 | 2.61 |
|  |  | 1Mb | 12.41 | 0.15 | 735.78 | 22.39 |
|  |  | 10Mb | 248.45 | 45.74 | 4346.00 | 260.88 |
|  | 500 | 100kb x 10 | 2.45 | 0.65 | 214.29 | 1.32 |
|  |  | 1Mb | 58.16 | 0.63 | 1363.26 | 89.77 |
|  |  | 10Mb | 458.04 | 40.77 | 5546.24 | 557.58 |
|  | 100 | 100kb x 10 | 36.47 | 0.38 | 359.09 | 3.49 |
|  |  | 1Mb | 483.45 | 69.74 | 3558.88 | 223.52 |
|  |  | 10Mb | 4394.58 | 172.11 | 29555.66 | 3257.71 |
| Coal | 1000 | 100kb x 10 | 0.48 | 0.02 | 128.45 | 3.78 |
|  |  | 1Mb | 7.55 | 2.15 | 461.96 | 51.90 |
|  |  | 10Mb | 95.63 | 3.01 | 2932.68 | 3.92 |
|  | 500 | 100kb x 10 | 1.41 | 0.12 | 196.38 | 19.48 |
|  |  | 1Mb | 18.86 | 0.53 | 936.67 | 26.22 |
|  |  | 10Mb | 127.02 | 7.45 | 3284.82 | 0.94 |
|  | 100 | 100kb x 10 | 16.48 | 1.49 | 307.23 | 14.25 |
|  |  | 1Mb | 104.91 | 23.49 | 2567.94 | 195.81 |
|  |  | 10Mb | 787.52 | 56.62 | 21024.12 | 1324.93 |
| Recap | 1000 | 100kb x 10 | 0.05 | 0.01 | 107.33 | 1.86 |
|  |  | 1Mb | 0.31 | 0.10 | 305.28 | 45.10 |
|  |  | 10Mb | 3.46 | 0.84 | 999.35 | 156.72 |
|  | 500 | 100kb x 10 | 0.13 | 0.02 | 172.39 | 2.41 |
|  |  | 1Mb | 1.23 | 0.37 | 806.97 | 94.56 |
|  |  | 10Mb | 7.53 | 2.11 | 1527.54 | 211.96 |
|  | 100 | 100kb x 10 | 2.06 | 0.46 | 244.49 | 2.93 |
|  |  | 1Mb | 24.75 | 7.33 | 2283.11 | 163.16 |
|  |  | 10Mb | 268.05 | 65.29 | 17097.95 | 1097.90 |

**Figure S1.**
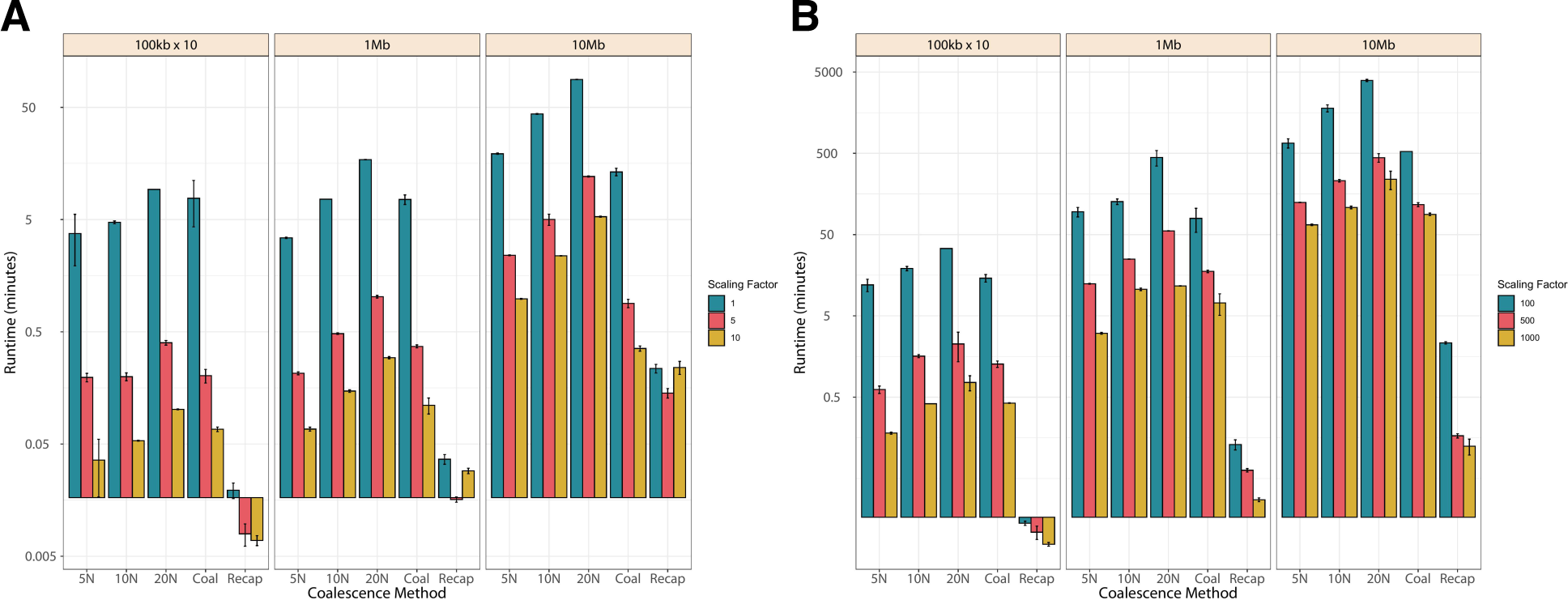
Runtime of the coalescent portion of the simulations human and *Drosophila* populations. Panel A) is the human data and Panel B) represents *Drosophila* data. The different colors represent different scaling factors, with blue being the lowest unscaled in human or 100 in *Drosophila* and yellow the highest 10 in human and 1000 in *Drosophila*. Each bar represents the average runtime of 10 simulation replicates with error bars showing standard deviation.

**Figure S2.**
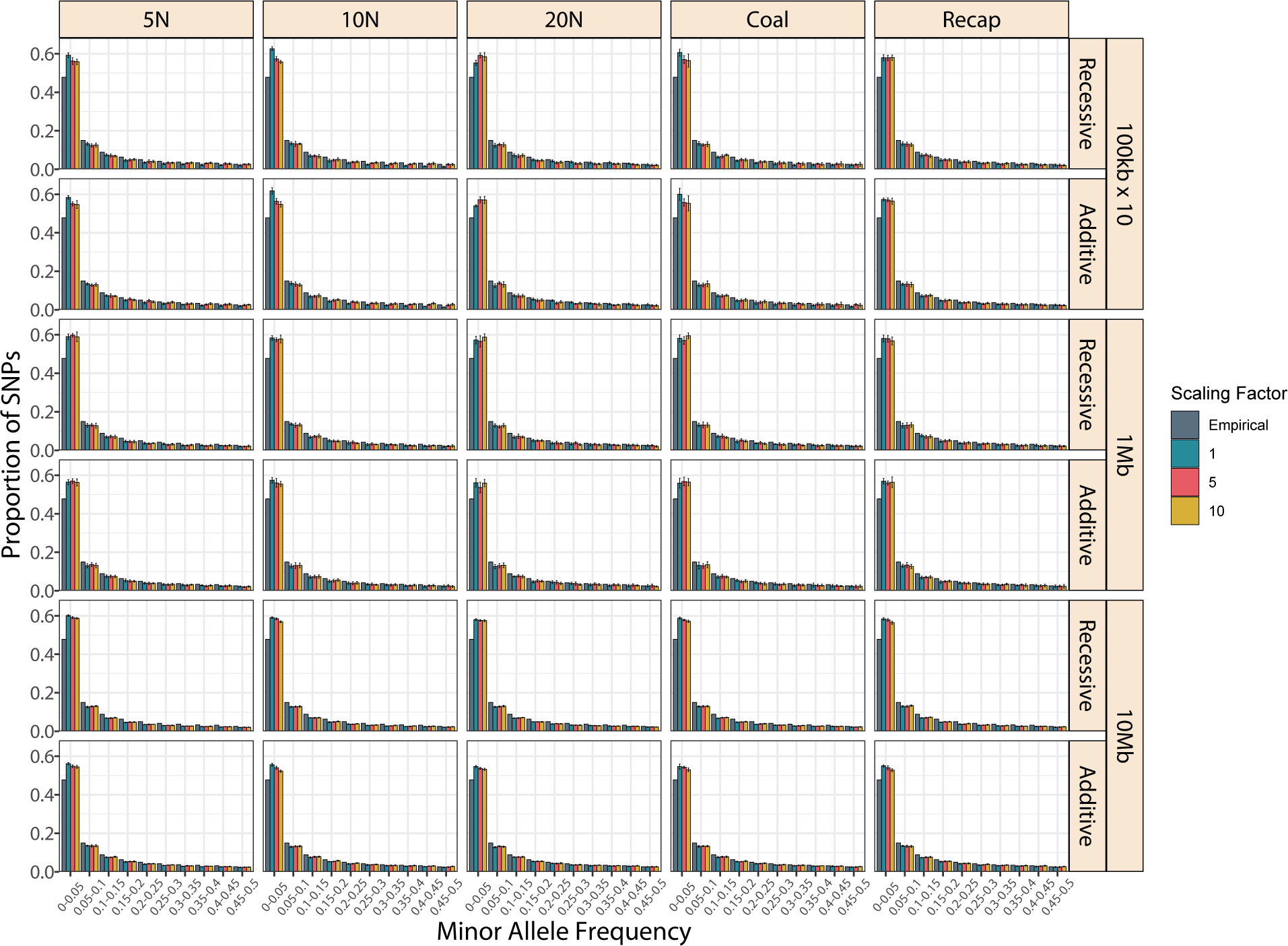
Whole genome site frequency spectra for simulated and empirical Yoruban population. Each row represents a different simulated segment length with an additive model for mutations and the situated above a recessive model of dominance. Different colors represent different scaling factors, with blue being the lowest unscaled and yellow the highest 10. MAF bins are left-exclusive and right-inclusive, meaning fixed sites (MAF=0) are not included.

**Figure S3.**
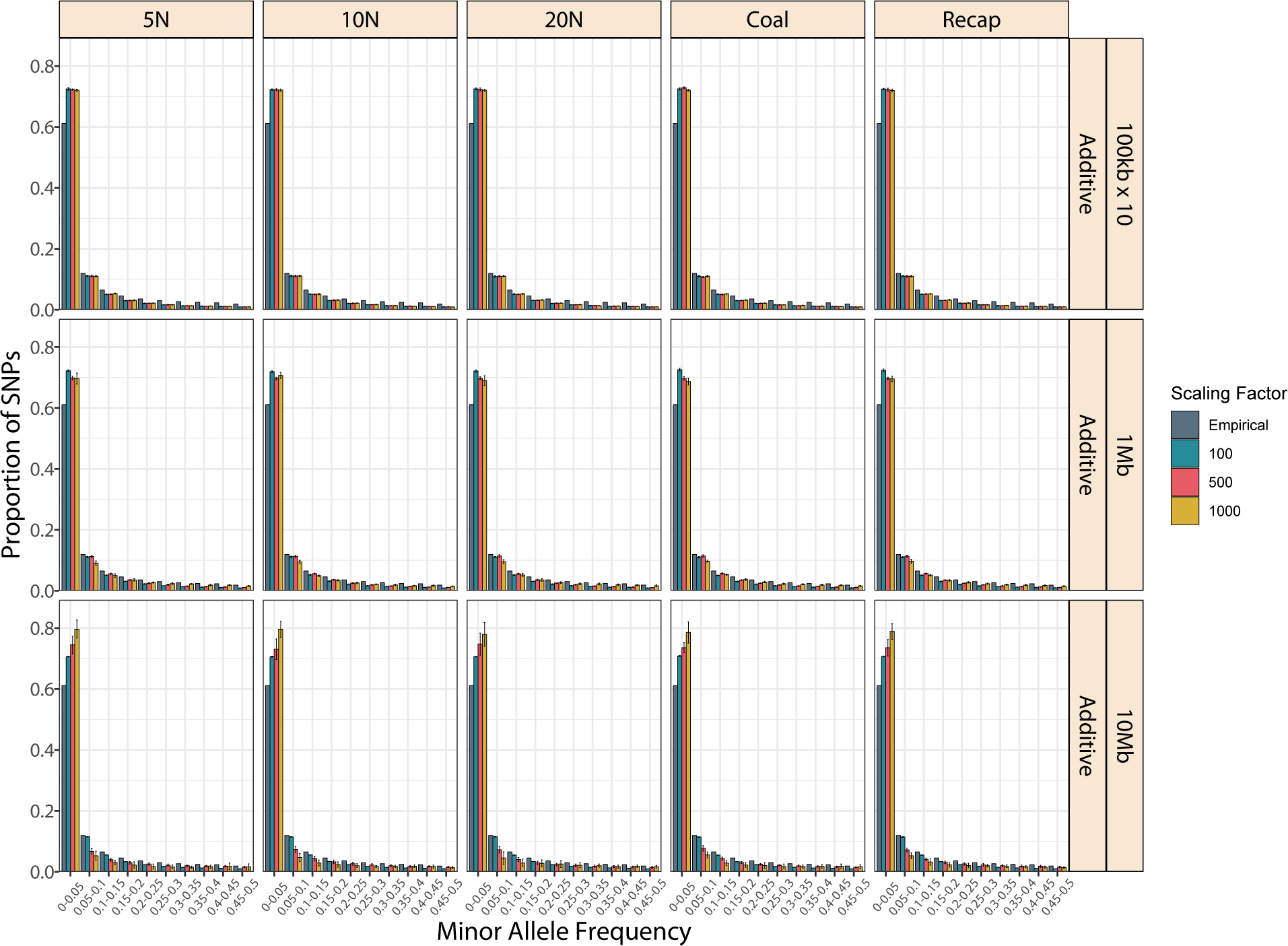
Whole genome site frequency spectra for simulated and empirical African *Drosophila* population. Different colors represent different scaling factors, with blue being the lowest 100 and yellow the highest 1000. MAF bins are left-exclusive and right-inclusive, meaning fixed sites (MAF=0) are not included.

**Figure S4.**
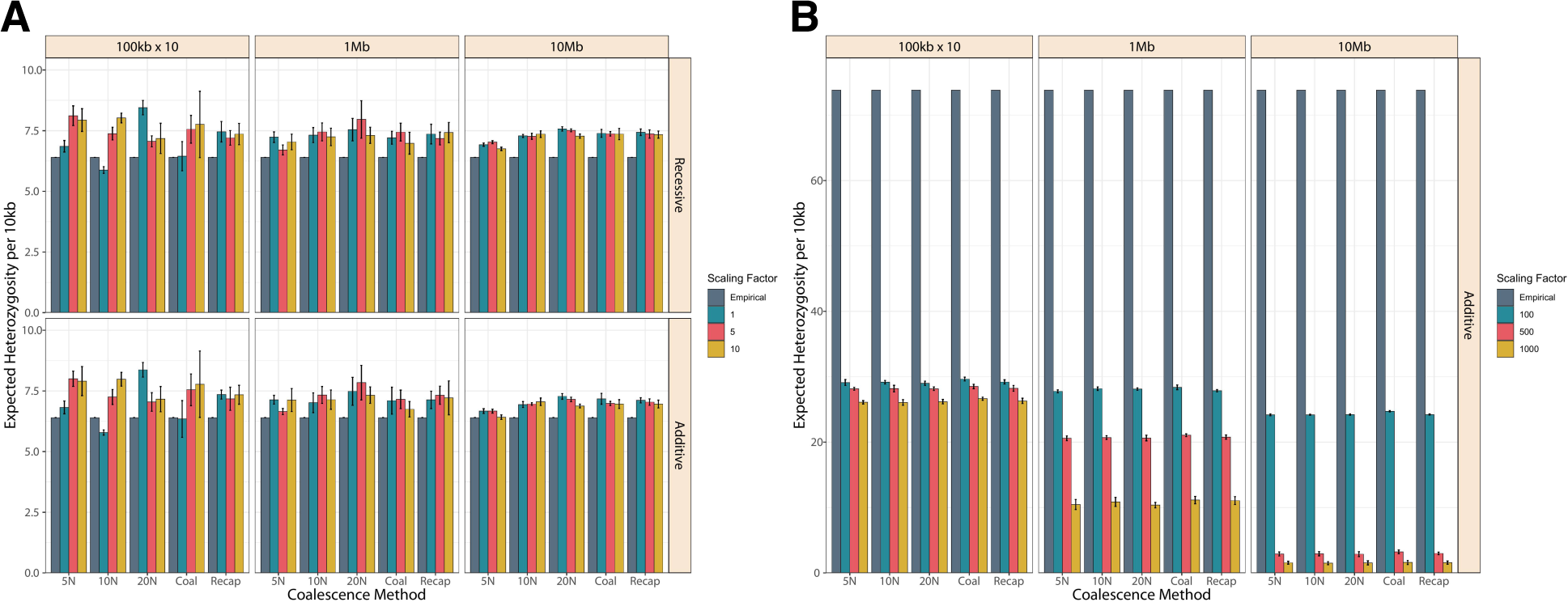
Heterozygosity computed from the whole genome site frequency spectra for simulated and empirical African populations. A) For human data, the top row used an additive model for mutations and the bottom row used a recessive model of dominance. Panel B) represents *Drosophila* data. The different colors represent different scaling factors, with blue being the lowest unscaled in human or 100 in *Drosophila* and yellow the highest 10 in human and 1000 in *Drosophila*. Each bar represents the average runtime of 10 simulation replicates with error bars showing standard deviation.

**Figure S5.**
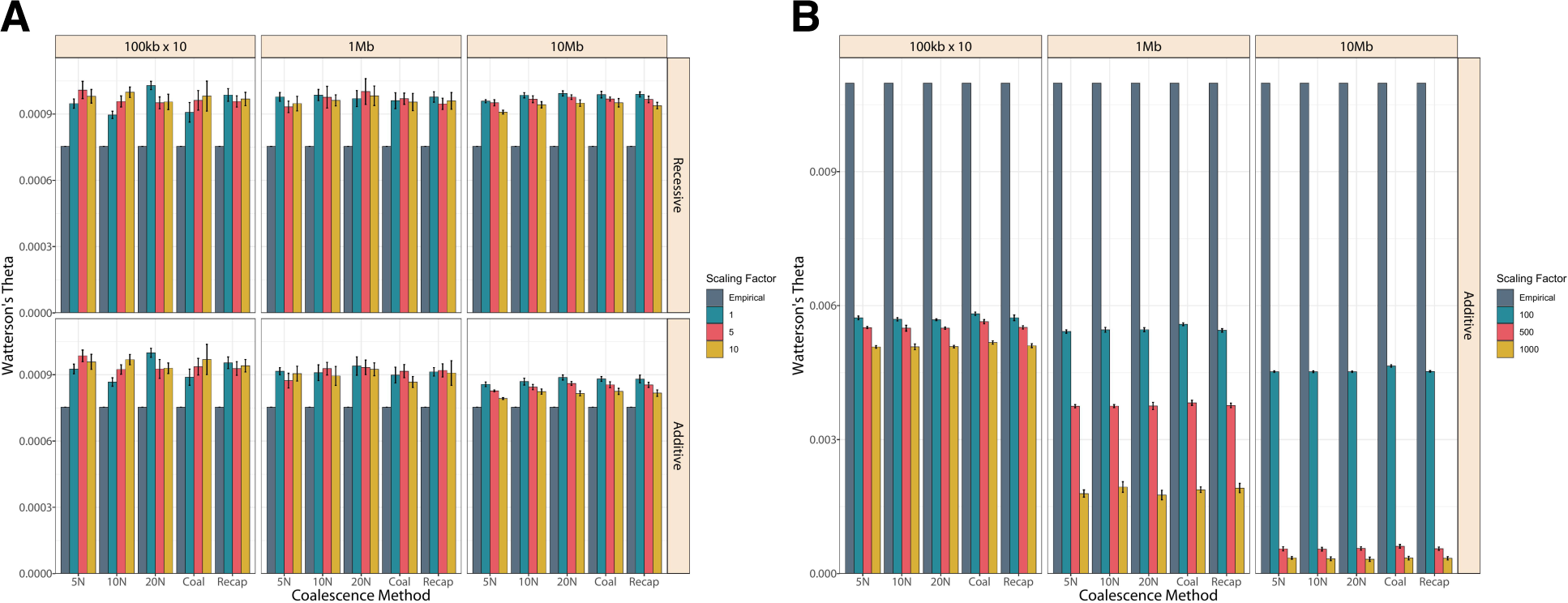
Watterson’s theta computed from the whole genome site frequency spectra for simulated and empirical African populations. A) For human data, the top row used an additive model for mutations and the bottom row used a recessive model of dominance. Panel B) represents *Drosophila* data. The different colors represent different scaling factors, with blue being the lowest unscaled in human or 100 in *Drosophila* and yellow the highest 10 in human and 1000 in *Drosophila*. Each bar represents the average runtime of 10 simulation replicates with error bars showing standard deviation.

**Figure S6.**
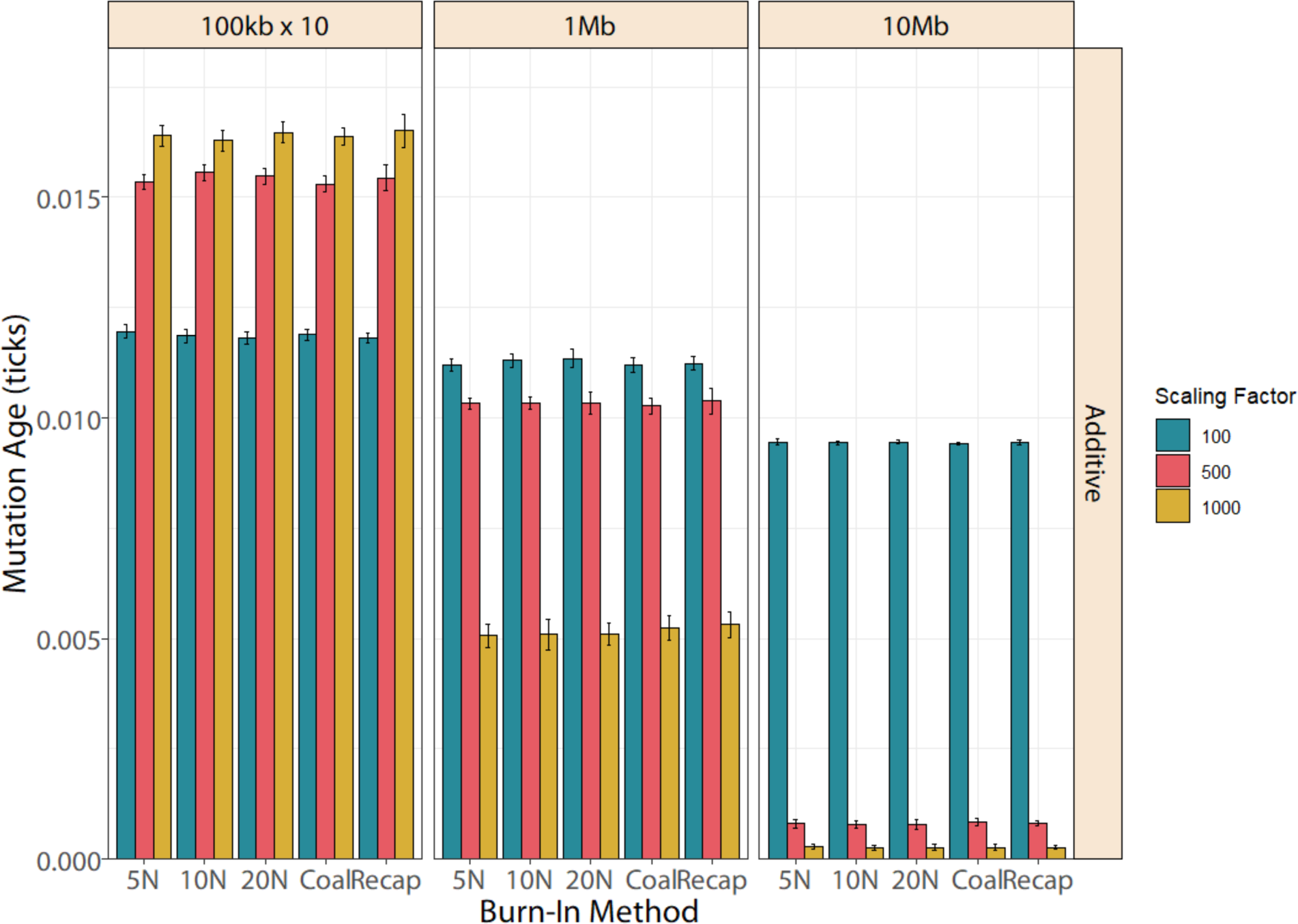
Mutation age across *Drosophila* simulations. Different colors represent different scaling factors, with blue being the lowest 100 and yellow the highest 1000.

